# A Structural Atlas of TAP Inhibition by Herpesviruses and Poxviruses

**DOI:** 10.1101/2025.06.19.660632

**Authors:** James Lee, Victor Manon, Jue Chen

**Author notes:** Corresponding author: Dr. Jue Chen (Contact author). These authors contributed equally (James Lee, Victor Manon).

## Abstract

In the host–pathogen arms race, herpesviruses and poxviruses encode proteins that sabotage the transporter associated with antigen processing (TAP), thereby suppressing MHC-I antigen presentation and enabling lifelong infection. Of the five known viral TAP inhibitors, only the herpes simplex virus protein ICP47 has been structurally resolved. We now report cryo-electron microscopy structures of TAP in complex with the remaining four: BNLF2a (Epstein–Barr virus), hUS6 (human cytomegalovirus), bUL49.5 (bovine herpesvirus 1), and CPXV012 (cowpox virus), assembling a structural atlas of viral TAP evasion. Employing divergent sequences, folds and conformational targets, these viral inhibitors converge on a common strategy: they stall TAP from the alternating access cycle, precluding peptide entry into the ER and shielding infected cells from cytotoxic T-cell surveillance. These findings reveal striking functional convergence and provide a structural framework for rational antiviral design.

## Introduction

Our immune system eliminates virus-infected cells by recognizing peptide antigens presented on the cell surface by major histocompatibility complex class I (MHC-I) molecules. These peptides are derived from intracellular proteins, transported into the endoplasmic reticulum (ER), loaded onto MHC-I, and exported to the cell surface. Foreign peptides—often originating from viral or tumor proteins—trigger an immune response that programs the affected cells for apoptosis^1,2^.

To evade this surveillance mechanism, many viruses have evolved strategies to disrupt antigen presentation^2–5^, often by targeting the transporter associated with antigen processing (TAP)—an ER-resident protein that delivers peptides to MHC-I^6^. TAP is an ATP-binding cassette (ABC) transporter composed of two homologous subunits, TAP1 and TAP2^7–11^. Each subunit contains an N-terminal transmembrane domain (TMD0) that interacts with other ER proteins, six transmembrane (TM) helices that form the peptide translocation pathway, and a nucleotide-binding domain (NBD) that binds and hydrolyzes ATP. The core TAP complex, lacking the TMD0s, is both necessary and sufficient for peptide transport^12–15^.

Like other ABC transporters^16,17^, TAP cycles between inward-and outward-facing conformations driven by ATP binding and hydrolysis^18–20^. In the inward-facing state, the NBDs are separated, and the peptide-binding cavity is open to the cytosol. Peptide and ATP binding trigger a conformational switch^19–22^ to the outward-facing state, where the NBDs dimerize and the transporter opens toward the ER lumen to release the peptide. ATP hydrolysis then resets TAP to its inward-facing state, initiating a new cycle of antigen transport.

Evidence that viruses directly target TAP to evade immune surveillance was first obtained with herpes simplex virus (HSV)^23–26^. It was noted that HSV-infected cells display markedly reduced MHC-I surface expression and are resistant to cytotoxic T-lymphocyte–mediated lysis^27,28^. The responsible molecule was soon identified as ICP47, an 88-residue cytosolic protein that binds TAP and blocks peptide translocation into the ER^23,24^. Subsequently, three independent studies uncovered US6, a structurally unrelated TAP antagonist encoded by several cytomegaloviruses (CMV)^29–31^. More recently, additional viral inhibitors—BNLF2a from Epstein–Barr virus (EBV)^32–34^, UL49.5 from several varicelloviruses^35,36^, and CPXV012 from cowpox virus (CPXV)^37–39^—have been shown to impair TAP-mediated peptide transport, underscoring the centrality of TAP as a viral immune-evasion target^4–6^.

Despite sharing a common function, these viral inhibitors display striking sequence and structural diversity. ICP47 is a soluble protein, while the other four are membrane-anchored with their functional domain either in the cytosol or the ER lumen (Fig. 1a)^4–6^. Despite comprehensive biochemical and cellular analyses of all five viral inhibitors, high-resolution structural data are available only for ICP47, which inserts an extended helical hairpin into the peptide-translocation pathway, occludes substrate binding, and locks TAP in its inward-facing conformation^40,41^.

**Fig. 1.**
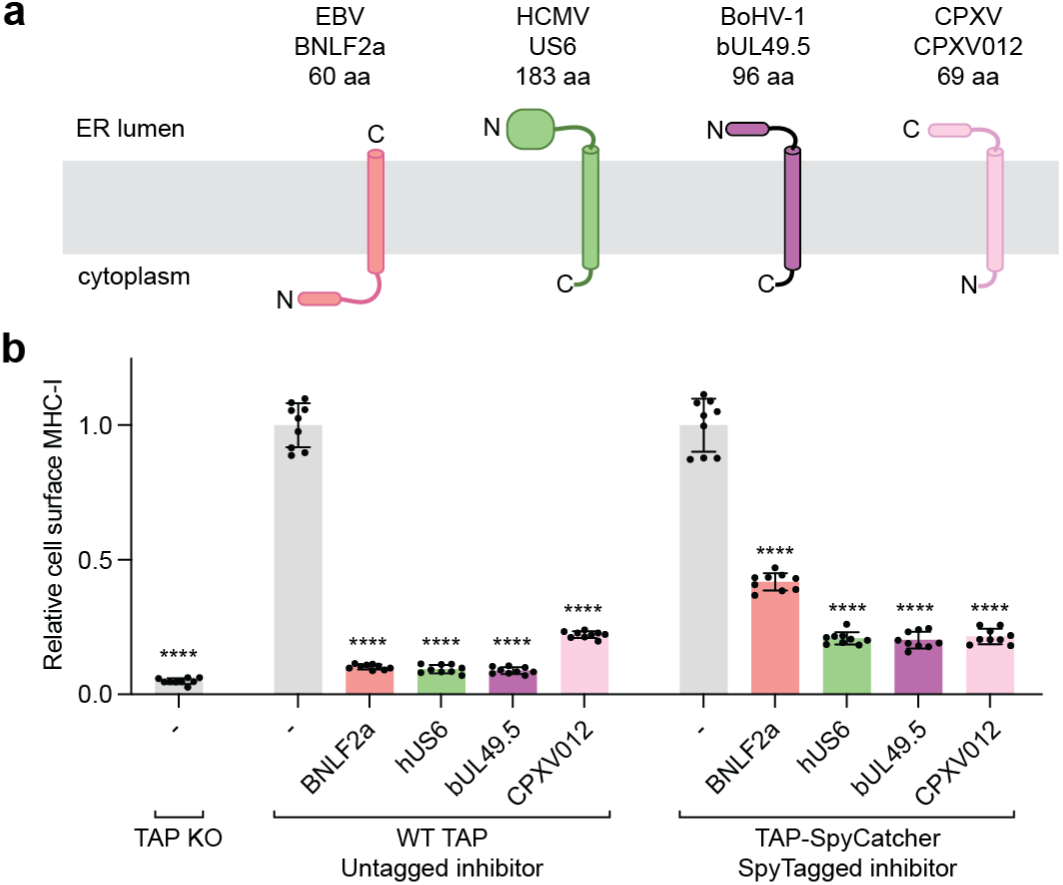
Viral inhibition of MHC-I antigen presentation. **a.** Topology diagram of the TAP viral inhibitors investigated in this study. The termini of the transmembrane helices are labeled. **b.** Functional assessment of the SpyCatcher/SpyTag system. Flow cytometry analysis of surface MHC-I levels in TAP-knockout cells transduced with baculoviruses expressing TAP and viral inhibitors. Data are normalized to the control cells without viral inhibitor and represent three technical replicates of three biological replicates (n=9). Error bars represent the standard error of the mean (SEM). Statistical significance relative to the (-) GFP only sample was tested by one-way analysis of variance. ****P<0.0001.

Elucidating the molecular structures of the remaining inhibitors is important for defining their binding sites and deciphering how each impairs TAP’s function.

In this study, we determined cryo-EM structures of TAP bound to each of the four remaining viral inhibitors: BNLF2a, US6, UL49.5, and CPXV012. These structures complete the molecular description of viral TAP inhibition and reveal a shared mechanism that has emerged through functional convergent evolution.

## Results

### A protein-tethering strategy to stabilize TAP/inhibitor complexes

To enable cryo-EM studies of TAP/viral inhibitor complexes, we developed a tethering strategy using the SpyCatcher/SpyTag system^42^. The C-terminus of the TAP1 NBD was fused to SpyCatcher, and the cytoplasmic terminus of each viral inhibitor was fused to SpyTag (Extended Fig. 1a). When the two modules are in proximity upon TAP/inhibitor binding, a covalent bond forms, preventing complex dissociation (Extended Fig. 1b). As expected, covalent TAP/inhibitor complexes formed efficiently in cells as evident by SDS-PAGE (Extended Fig. 1c) and fluorescence size exclusion chromatography (Extended Fig. 1d-e).

We evaluated whether the tagged TAP and inhibitors remained functional and could assemble into complexes in cells by flow cytometry (Extended Fig. 1f). TAP-knockout cells display few MHC-I molecules on the cell surface (Fig. 1b). Expression of the TAP-SpyCatcher construct restored surface MHC-I to a level comparable to that of the wild-type (WT) TAP (Extended Fig. 1g). Tagged viral proteins inhibited both WT and the SpyCatcher-tagged TAP (Fig. 1b and Extended Fig. 1h); and except for BNLF2a, the activities of the tagged proteins were similar to those of their untagged counterparts (Fig. 1b). A potential explanation for the slight decrease in activity observed with tagged BNLF2a will be discussed in the context of its structure.

### Epstein-Barr virus protein BNLF2a traps TAP in the inward-facing conformation

EBV is a gammaherpesvirus that establishes persistent infection in >90% of the human adult population^43^. Although EBV infections acquired in childhood are typically asymptomatic, infections in adults can cause mononucleosis and are linked to cancer risk^43^. EBV encodes several proteins to downregulate antigen presentation, one of which is the TAP inhibitor BNLF2a^44–46,33^. BNLF2a is a tail-anchored protein with an N-terminal cytosolic domain and a single-pass, C-terminal transmembrane (TM) helix^32–34^ (Fig. 1a). Homologs of BNLF2a are only found across Old World primate gammaherpesviruses, and all can inhibit human TAP^33^. Functional experiments suggest that BNLF2a interacts directly with core TAP and that both the cytosolic and transmembrane domains of the inhibitor are necessary for inhibition^33,47,48^.

The structure of the TAP/BNLF2a complex was determined at an overall resolution of 3.7 Å in the absence of nucleotide (Fig. 2 and Extended Fig. 2). TAP adopts an inward-facing, NBD-separated conformation (Fig. 2a) closely resembling that of the apo TAP structure (PDB code 8T46), with an overall root mean square deviation (RMSD) of 0.96 Å. Clear density corresponding to BNLF2a allowed us to model all but five residues of the viral inhibitor (Extended Fig. 2c-d). BNLF2a is shaped like a fishing hook, with residues 5-27 forming a hairpin structure, traversing ∼20 Å across the TM cavity before looping back (Fig. 2b). A proline-rich region (residues 28–36) forms the “bend”, exiting the TM cavity through the cytoplasmic opening between TAP2 TM helices 4 and 6 (Fig. 2b-c). The C-terminal TM helix of BNLF2a represents the “bend,” packing along the external surface of TAP2 and extends into the ER lumen (Fig. 2a-c).

**Fig. 2.**
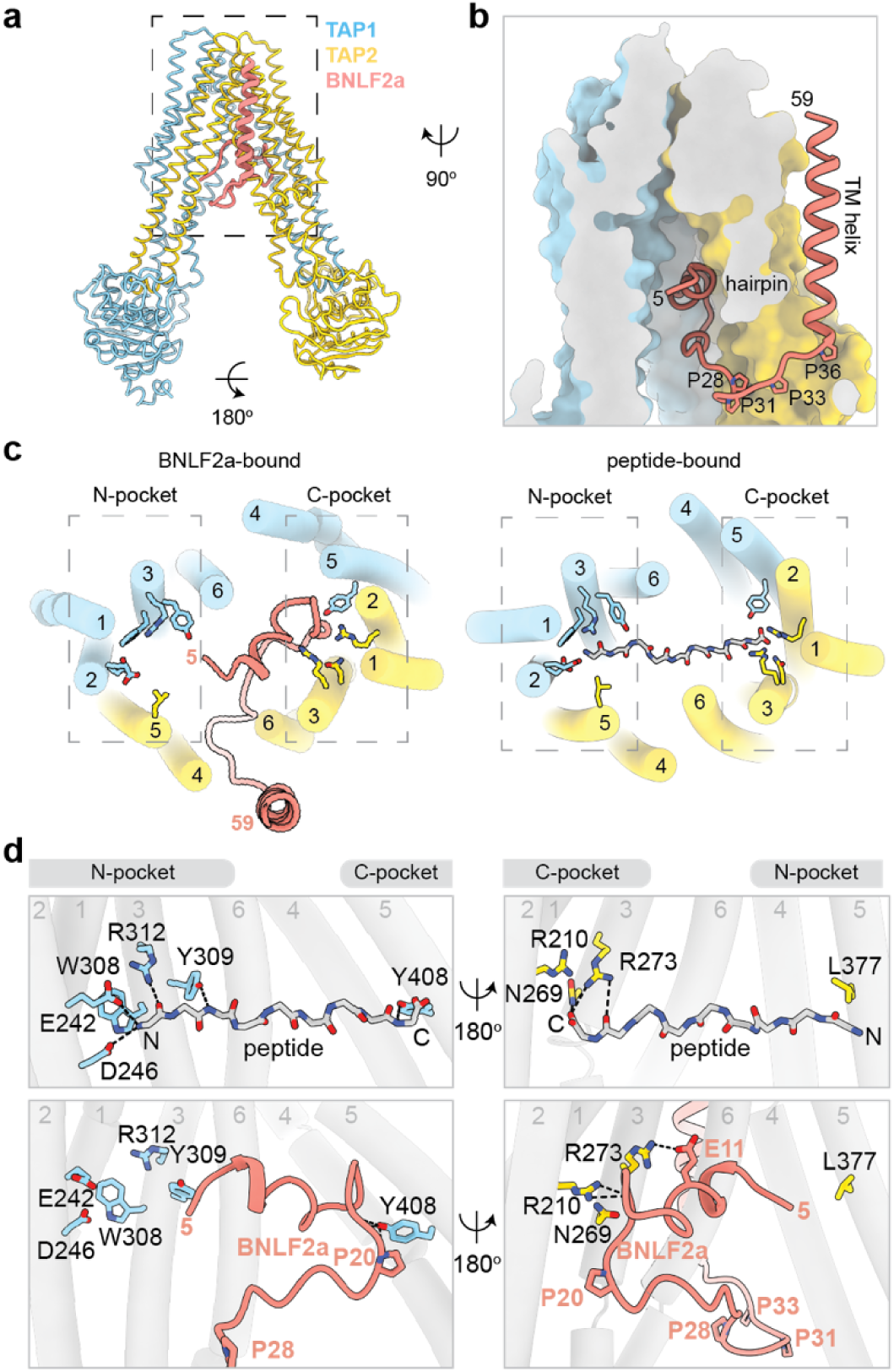
BNLF2a from Epstein-Barr virus inserts an N-terminal helix from the cytoplasm into inward-facing TAP. **a.** The overall structure. TAP1, TAP2, and BNLF2a are colored in sky blue, gold, and salmon, respectively. **b** Zoomed-in view of the TAP translocation pathway sliced perpendicular to the plane of the membrane. TAP is shown as molecular surface. The first and last visible BNLF2a residues are numbered. **c.** Left: BNLF2a interacts with the C-pocket of the TAP substrate binding site, from the ER lumen. Right: The TAP N-and C-pockets in the inward-facing state interact with a peptide antigen (PDB: 8T4F). **d.** Details of TAP/ peptide (upper) and TAP/BNLF2a (bottom) interactions at the N-and C-pockets. Pocket residues in TAP1 are shown in blue on the left panel and those of TAP2 in yellow on the right panel. Hydrogen bonds are shown as dotted lines.

Biochemical data have shown that BNLF2a inhibits peptide binding^33,47^, and the structure shows that it does so by occupying the same space as the peptide antigen (Fig. 2c-d). TAP recruits peptides from the cytosol in the inward-facing, NBD-separated conformation^49^. Peptides of diverse sequences bind in a similar manner, anchoring their N-and C-terminal ends into two distinct sites on TAP, named the N-and C-pockets (Fig. 2c-d). The hairpin structure of BNLF2a inserts into the peptide-binding cavity and completely obstructs the C-pocket through a network of main-chain and side-chain interactions (Fig. 2c-d). In the structure, the first ordered residue of BNFL2a, a leucine in position 5, is positioned 10Å from the N-pocket of TAP (Fig. 2d). Given that the tethering strategy introduced a SpyTag at the N-terminus of BNLF2a (Extended Fig. 1), it remains possible that in its native, untethered form, BNLF2a engages the N-pocket via its free N-terminus, mimicking a peptide antigen. Indeed, modelling of the untagged BNLF2a structure places the N-terminus of BNLF2a within the amino-termini accepting site of the N-pocket (Extended Fig. 2h). Such an interaction could explain the observed reduction in inhibitory activity of the tethered construct compared to free BNLF2a (Fig. 1b).

These structural observations indicate that, similar to ICP47^23–26^, the cytosolic domain of BNLF2a acts as a competitive inhibitor of the peptide antigen^33,47^. However, unlike ICP47, BNLF2a requires a TM anchor, as the cytoplasmic domain alone (ΔTM) was insufficient to inhibit TAP^47,48^ (Extended Fig. 2g). This difference may be explained by the smaller interface between BNLF2a and TAP: the solvent-accessible surface area of TAP buried by BNLF2a is 1387 Å², approximately 60% that of ICP47 (2360 Å²). Specific interactions with TAP within the membrane are not essential, since chimeric BNLF2a constructs containing a synthetic polyleucine helix or the TM helix from another tail-anchored protein^48^ showed no defect in TAP inhibition (Extended Fig. 2g). These results suggest that the TM anchor primarily functions to enrich BNLF2a in the ER membrane, thereby increasing the local concentration of its inhibitory cytoplasmic domain.

In addition to competing directly with peptide binding, BNLF2a also prevents TAP from transitioning to the outward-facing conformation, which involves closure of the cytoplasmic opening between TM4 and TM6 upon NBD dimerization. In the TAP/BNLF2a complex, this conformational change is blocked because the proline-rich bend wedges between TAP2 TM4 and TM6 (Fig. 2b-c), effectively jamming TAP in an inward-facing state. Therefore, BNLF2a inhibits TAP by simultaneously occupying the peptide-binding site and blocking conformational changes required for peptide transport.

### US6 of cytomegalovirus traps TAP in an NBD-dimerized, post-hydrolytic conformation

Like EBV, human cytomegalovirus (HCMV) is a clinically significant herpesvirus that establishes persistent infections in human hosts. While acute infections present with mild disease in healthy hosts, latent infection is associated increased cancer risk and can be threatening in those with immature or compromised immune systems^29,30^. US6 from CMV is a type I TM protein consisting of a large luminal region, a TM helix, and a short cytosolic tail (Fig. 1a). Mutagenesis studies have shown that, unlike ICP47 and BNLF2a, the inhibitory domain of US6 lies entirely within its ER luminal region^29,30^, suggesting that US6 operates through a mechanism distinct from those of ICP47 and BNLF2a. Interestingly, despite significant sequence divergence, human US6 (hUS6) and rhesus CMV US6 (rhUS6 or RH185) both inhibit human TAP^50^. To gain structural insights into the molecular basis for US6-mediated inhibition, we determined the cryo-EM structures of TAP bound to either hUS6 or rhUS6, revealing a conserved mode of action despite their limited sequence homology (Fig. 3 and Extended Fig. 3).

**Fig. 3.**
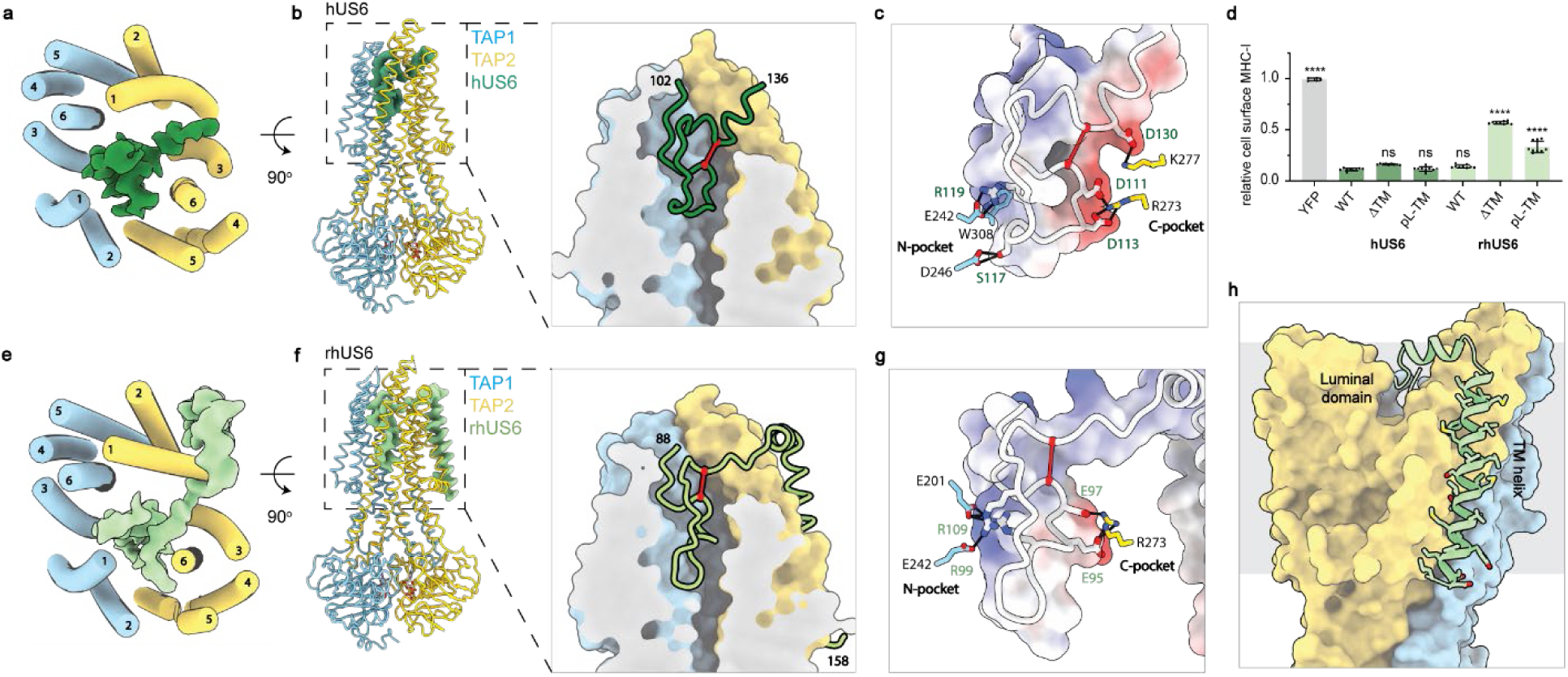
US6 from cytomegalovirus inserts a disulfide-bonded loop into outward-facing TAP. a-b: Structure of the TAP/hUS6 complex. TAP is colored as in Fig. 2 and the cryo-EM density for hUS6 is shown in green. ATP is shown as sticks. (Right) Zoom-in view of the TAP translocation pathway. TAP is shown as a molecular surface and hUS6 in ribbon. hUS6 cysteines 110 and 129 are denoted as red balls and connected via red line. The first and last hUS6 residues observed in the map are numbered. **c.** Electrostatic properties of hUS6 complements those of the N-and C-pockets. hUS6 is represented as white wire superimposed onto a molecular surface colored by electrostatic potential. Scale: from-10 kT/e (red) to +10 kT/e (blue). Interacting residues between hUS6 and TAP N-and C-pockets are shown and labelled. **d.** Effect of US6 TM variants on cell-surface MHC-I assays. Data represent the relative MHC-I cell surface levels normalized to the control (YFP only) and represent three technical replicates of three biological replicates (n=9). Error bars represent the SEM. Labels: wild-type (WT), poly-leucine TM helix (pL-TM), TM deletion (ΔTM). Statistical significance relative to treatment with wild-type hUS6 IRES YFP was tested by one-way analysis of variance. ****PP<0.0001; ns > 0.05. **e-f.** Structure of the TAP/rhUS6 complex. **g.** Electrostatic properties of rhUS6 and the interactions with N-and C-pocket residues, as scale as in panel c. **h.** TM helix of rhUS6 packs along TAP2. TAP and rhUS6 are shown as molecular surface and ribbons, respectively.

Both structures were determined with WT TAP in the presence of 10 mM ATP to approximately 3 Å resolution. In both cases, TAP exhibits an NBD-dimerized, outward-facing conformation, but its luminal cavity is completely plugged by the viral inhibitor (Fig. 3). For hUS6, this plug consists of residues 102–136 of its luminal region, which forms a large loop stabilized by intramolecular packing and a disulfide bond between two conserved cysteines (Fig. 3a-b). Residues outside this region were invisible in the cryo-EM map, suggesting they are highly flexible. The structured region closely aligns with previous studies that pinpointed the functional domain of hUS6 to lie between 98-143^51^. Mutating or removing residues outside this region did not affect MHC-I downregulation^29,30,51^. Furthermore, tetra-alanine scanning studies identified residues 107–118 as particularly critical for TAP inhibition^51^. This region extends deep into the transmembrane (TM) cavity, interacting with TAP residues that are highly conserved for binding peptide antigens (Fig. 3c). In particular, hUS6 residues S117 and R119 insert into the negatively charged N-pocket while two acidic residues, D111 and D113 interact with the basic residues in the C-pocket (Fig. 3c). These interactions confer the high affinity of hUS6, enabling TAP inhibition even in the absence of the TM anchor or the N-terminal region^29,30,51^ (Fig. 3d).

The structure of the TAP/rhUS6 complex also reveals a disulfide-tethered loop that plugs the ER-facing lumen of TAP (Fig. 3e-g). Like its homolog in human CMV, rhUS6 also interacts with residues in the N-and C-pockets through electrostatic interactions (Fig. 3g). Compared to hUS6, the main-chain atoms follow a different trajectory and descend deeper into the TM cavity though the overall shape and electrostatic distribution remain very similar (Fig. 3c,g). A unique feature of rhUS6 is the well-ordered TM helix, which packs tightly against the exterior surface of TAP2 (Fig. 3h). While truncating or replacing the TM helix of hUS6 had no functional consequence— consistent with previous studies^29,52^—similar modifications in rhUS6 severely impaired its inhibitory function (Fig. 3d). Thus, whereas the TM helix of hUS6 is structurally unresolved and dispensable for function, the ordered TM helix of rhUS6 is crucial for effective TAP inhibition, underscoring a clear structure-function relationship.

Finally, previous studies suggested that hUS6 prevents TAP from binding ATP^52,53^. However, our cryo-EM reconstructions of both hUS6-and rhUS6-bound TAP show densities corresponding to ATP bound at the degenerate site and ADP at the consensus site (Extended Fig. 3e,i). These observations indicate that US6 does not directly block nucleotide binding *per se*. Instead, US6 binding traps TAP in an NBD-dimerized, post-hydrolytic state, thereby preventing nucleotide exchange and subsequent ATP hydrolysis cycles required for peptide transport. This mechanism effectively halts TAP activity by locking the transporter into an inactive conformation, rather than by inhibiting ATP binding itself.

### UL49.5 from bovine herpesvirus 1 is a dual-action immune evasion protein

Although bovine herpesvirus 1 is not a human pathogen, it encodes bUL49.5, a type I TM protein that inhibits TAP across different species including human^54,55^. bUL49.5 is unique among the known viral inhibitors due to its dual action as an inhibitor of peptide transport and also for targeting TAP for proteasomal degradation^35,36,54–56^. Studies have mapped these two functions to distinct regions: the N-terminal ER-luminal domain and TM inhibit peptide transport, while the C-terminal cytosolic tail encodes a degron that marks TAP for ERAD-mediated degradation^35,36,55^ (Fig. 1a). C-terminal inhibitor truncations fail to degrade TAP but still bind and inhibit the transporter, signifying that bUL49.5 is a bona fide TAP inhibitor^35,36,55^.

From a single cryo-EM dataset collected with the TAP E632Q (EQ) variant incubated with 10mM ATP, we observed two structures of the TAP/bUL49.5 complex (Fig. 4 and Extended Fig. 4). Both show an NBD-dimerized TAP with its ER-facing cavity plugged by the viral inhibitor (Fig. 4a-b). The two structures differ in the local conformation of TAP2 TM3: in the “unkinked” conformation, TM3 is continuous whereas in the “kinked” conformation, TM3 exhibits a sharp inward kinking that is stabilized by the presence of the inhibitor and interactions between R467 of TAP1 and N267 of TAP2 (Fig. 4c).

**Fig. 4.**
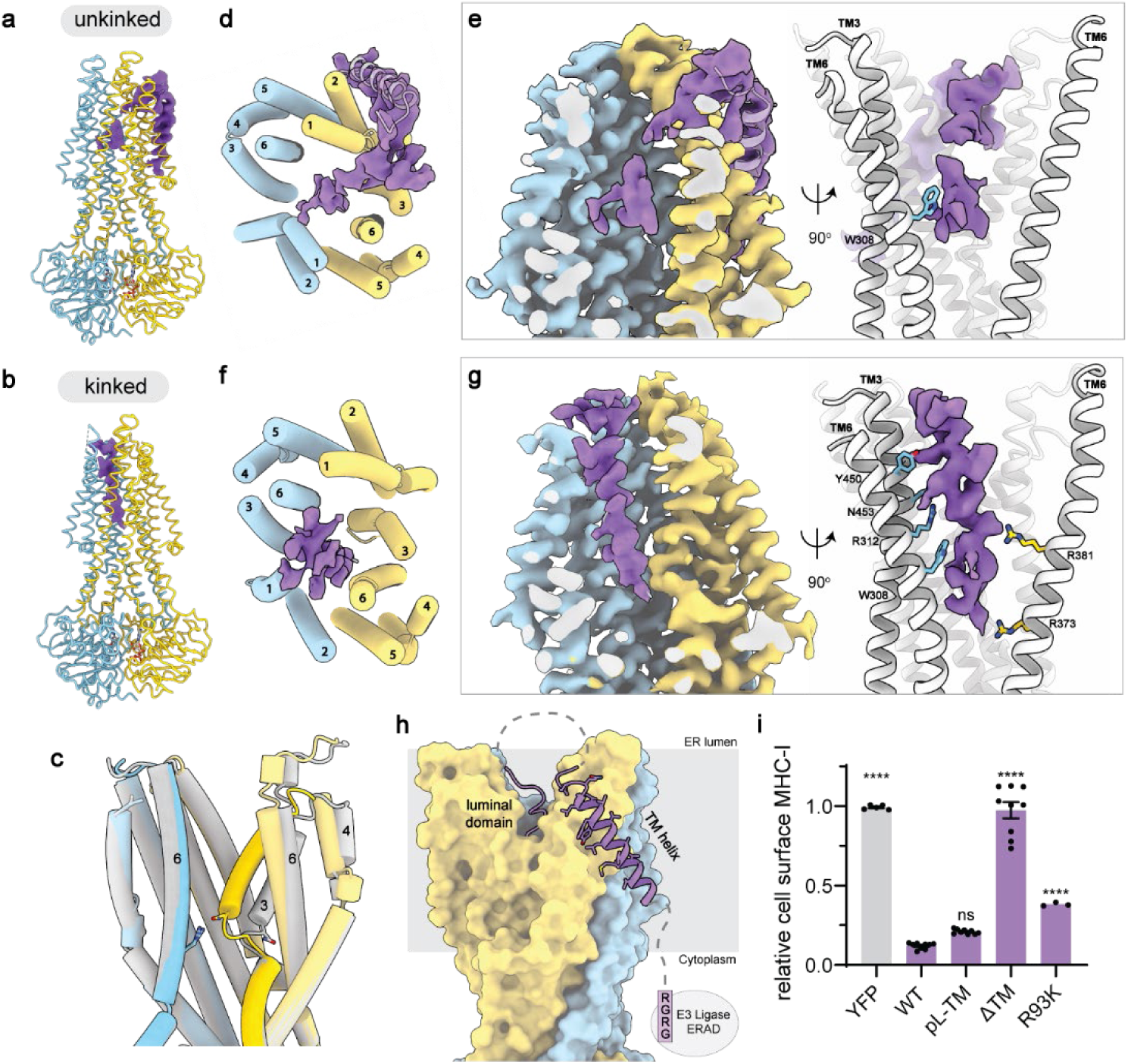
bUL49.5 from bovine herpes virus 1 inserts its N-terminal luminal region into outward-facing TAP. **a,b** TAP adopts two conformations in the presence of bUL49.5. The cryo-EM density for bUL49.5 in both conformations is shown in purple. **c.** Superposition of the two TAP structures, illustrating conformational differences within the translocation pathway. The kinked structure is shown in color and unkinked structure in grey. The side chains for R467 of TAP1 and N267 of TAP2 are shown as sticks. **d.** The unkinked TAP/bUL49.5 complex viewed from the ER-lumen, also shown in grey ribbon is the TM helix of bUL49.5. **e.** (Left) Cross-section of the plugged TAP translocation pathway in the unkinked conformation. Densities corresponding to TAP are in blue/yellow, and inhibitor in purple. (Right) Rotated view with the side chain of TAP1 W308 illustrated. **f.** The kinked TAP/bUL49.5 structure. **g.** Density (Left) and contact residues (Right) of the kinked structure. **h.** Topology of bUL49.5, TAP (unkinked) is shown as a molecular surface. Sequences not visible in the density are represented as dashed lines. The C-terminal RGRG degron (not observed in the map) is indicated. **i.** Effect of bUL49.5 TM variants and the degron deficient mutation, R93K, on cell-surface MHC-I assays, normalized to the YFP only sample. Data for WT, pL-TM, and ΔTM samples represent three technical replicates of three biological replicates (n=9). R93K data points represent one biological replicate in technical triplicate (n=3). Error bars represent the SEM. Statistical significance relative to treatment with wild-type bUL49.5 IRES YFP was tested by one-way analysis of variance. ****PP<0.0001; ns > 0.05.

In the more populated “unkinked” conformation (56% of total particles), densities were well-defined for the side chains in TAP and the two ATP molecules at the NBD dimer interface (Extended Fig. 4f). However, the density for bUL49.5 is discontinuous, with discernible features limited to the TM helix and the luminal domain (Fig. 4d-e). Like US6, the N-terminal luminal region of bUL49.5 inserts into the TM cavity, penetrating more than 20 Å into the lipid bilayer. Despite the highly heterogeneous and discontinuous density of bUL49.5, TAP1 W308 in the N-pocket forms a clear contact with the inhibitor (Fig. 4e, right).

In the less-populated “kinked” conformation (27% of total particles), TAP is similarly trapped in an ER-facing state (Fig. 4b,f,g). Compared to the “unkinked” complex, the density for bUL49.5 inside the TM cavity is better defined, though still insufficient to assign the amino acid register (Fig. 4g). The improved ordering of the inhibitor density likely results from a larger contact area with TAP. In addition to W308, several other residues in TAP also interact with the inhibitor, including N-pocket residue R312 (Fig. 4g, right).

The TM helix of bUL49.5 was only resolved in the “unkinked” reconstruction, which packs diagonally across the exterior surface of TAP2 (Fig. 4h). No density was observed for the C-terminal degron sequence in either map. Mutational studies have shown that the TM helix of bUL49.5 is essential for MHC-I downregulation, but its specific sequence is not critical^55^. Consistent with this conclusion, we found that deleting the TM helix abolishes its inhibitory function, whereas replacing it with a poly-leucine helix still potently inhibits TAP (Fig. 4i). We also observed that ablating the C-terminal degron via an R93K substitution weakens bUL49.5 inhibition^55,57^, indicating that promoting TAP degradation is an important component of the mechanism by which bUL49.5 downregulates MHC-I surface expression (Fig. 4i).

Altogether, these results suggest that bUL49.5 disrupts peptide transport through the concerted action of a low-affinity TAP-binding domain that both inhibits the transporter and marks it for degradation via a cytoplasmic degron. The amorphous density for the inhibitor suggests that the specific interactions between bUL49.5’s luminal domain and TAP are low affinity, rendering it a weaker inhibitor. However, this weaker inhibitory function is compensated by the cytoplasmic degradation domain, which targets TAP for ERAD-mediated degradation. Through these coordinated mechanisms—direct inhibition of peptide transport and degradation of TAP— bUL49.5 ensures efficient suppression of antigen presentation, ultimately facilitating immune evasion.

### CPXV012, a TAP inhibitor from the cowpox virus

Outside the *Herpesviridae* family, the cowpox virus of the *Poxviridae* family encodes several immune-evasion proteins including the TAP inhibitor CPXV012^38,58^. Infection with the cowpox virus, although largely benign in healthy humans, can be lethal in immunocompromised patients^59,60^. The 69-residue protein CPXV012 comprises an N-terminal TM helix and a C-terminal ER-luminal domain (Fig. 1a). Coimmunoprecipitation assays have demonstrated that CPXV012 directly interacts with TAP^61^; however, its precise functional domain and mechanism of inhibition remain poorly defined. One model suggests that the TM helix facilitates binding to TAP, while the C-terminal ten residues inhibit by mimicking a high-affinity peptide^61^. In contrast, an alternative study proposes that the TM helix is largely dispensable and that the interaction of the luminal domain with the lipid membrane is essential^62^. These different views highlight the need for further structural and functional studies.

The structure of the TAP(EQ)/CPXV012 was determined in the presence of 10mM ATP to 3.0 Å resolution (Fig. 5 and Extended Fig. 5). Like US6 and bUL49.5, the viral inhibitor stabilizes TAP in an outward-facing conformation via its ER luminal domain (Fig. 5a), which in this case forms a well-defined helix inserting approximately 25 Å into the TM cavity of TAP (Fig. 5b). No density was observed for the N-terminal 40 residues (Extended Fig. 5b-d), suggesting they are flexible and do not form stable interactions with TAP. The helix, consisting of residues 45–61, packs tightly against TAP1 TM1, 3, 6, and TAP2 TM1, 2, 4 (Fig. 5c-d). Following a sharp bend (Fig. 5b), the C-terminal seven residues of CPXV012 stretch across the TM cavity and reach the C-pocket (Fig. 5e). The terminal carboxyl group of the viral inhibitor forms two hydrogen bonds with C-pocket residues, TAP2 N269 and R273^49^ (Fig. 5f and Extended Fig. 5i). The N-pocket is also occupied by the viral inhibitor (Fig. 5f), with the side chain R59 mimicking the free amino group of the peptide to form a cation-π interaction with TAP1 W308 and a hydrogen bond with TAP1 E242 (Fig. 5f and Extended Fig. 5h).

**Fig. 5.**
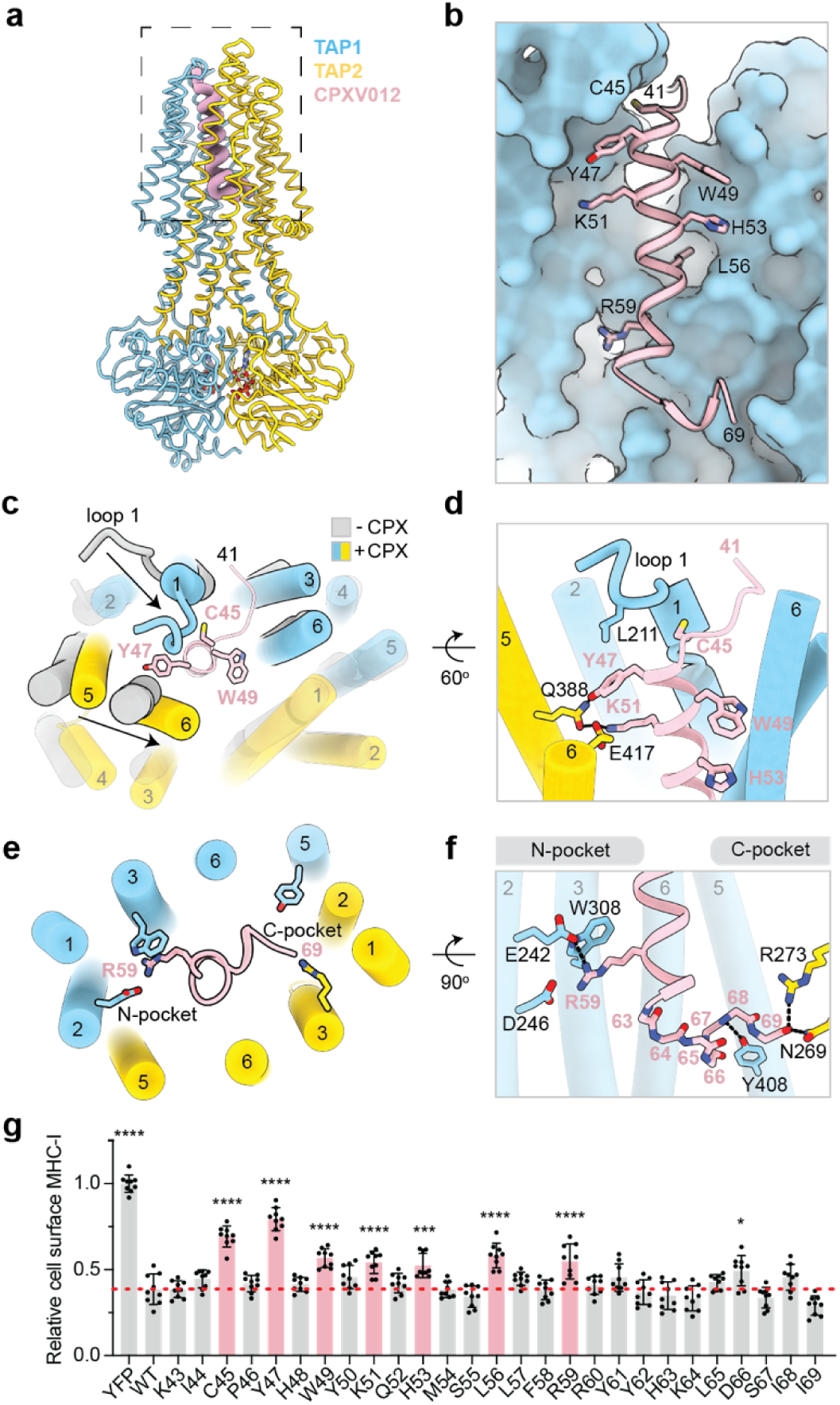
CPXV012 from cowpox virus inserts a C-terminal helix from the ER lumen into outward-facing TAP. **a.** Molecular model of TAP(EQ) bound to CPXV012 (pink wire). ATP is shown as sticks and colored by heteroatom. **b.** Zoom-in of the TAP translocation pathway. TAP1 is shown as surface and CPXV012 is shown in ribbon. Residues identified in **g** are shown as sticks. **c.** Superposition of the structures of outward-facing TAP in the absence (grey) or presence (colored) of CPXV012. The conformational changes upon binding of CPXV012 are highlighted. **d.** Zoom-in of the interface between TAP and CPXV012 at the ER-luminal entrance of the TM cavity. **e.** Zoom-in of the interface near the N-and C-pockets. **f.** Interactions of CPXV012 with the TAP N-and C-pockets. The main chain of the last seven residues of CPXV012 are represented as sticks and numbered by their alpha carbons. **g.** Effect of single alanine substitutions in the ER luminal domain of CPXV012. Flow cytometry analysis of HEK293S cells expressing CPXV012 single alanine substitution variants coupled to YFP with an IRES element. Substitutions that are statistically significant are colored pink and shown in panel **b**. The red dotted line marks the relative cell surface MHC-I levels in the presence of WT CPXV012. Data represent the relative MHC-I cell surface levels normalized to the YFP only sample and represent three technical replicates of three biological replicates (n=9). Error bars represent the SEM. Statistical significance relative to treatment with wild-type CPXV012 IRES YFP was tested by one-way analysis of variance. ****PP<0.0001; ***P(H53A)=0.005,*P(D66A)=0.0152. All other pairwise comparisons to the wild-type were not significant.

Compared to the inhibitor-free structure, conformational changes upon CPXV012 binding occur locally in the ER-luminal half of the TMDs (Fig. 5c). The largest change is in the luminal loop following TAP1 TM1, which swings ∼45° toward the inhibitor (Fig. 5c-d). Additionally, TAP2 TM5 and TM6 move inward to make direct contact with the helix (Fig. 5c-d). Consequently, the TM cavity, which opens the peptide-binding site to the ER lumen, is now obstructed by the viral inhibitor.

The interface between TAP and the viral inhibitor is extensive (2106 Å^2^) and single alanine substitution assays indicated that several residues are critical for inhibition (Fig. 5g). Among them, C45, located at the top of the helix, interacts with the TAP1 luminal loop 1 (Fig. 5d). Y47 and K51, positioned on the same face of the helical handle, form a hydrogen bonding network with TAP2 Q388 and E417 (Fig. 5d). These residues are directly involved in stabilizing the conformational changes induced upon inhibitor binding (Fig. 5c-d). Additionally, interactions with the N-pocket, mediated by R59, and contacts with TAP1, mediated by W49, H53, and L56, are also important (Fig. 5f). Alanine substitution of any of these residues significantly reduces CPXV012 inhibition (Fig. 5g), supporting the model that the entire luminal domain—not just the C-terminal 10 residues—is crucial for inhibition.

Although we did not observe EM density for the TM helix, MHC-I presentation assays reveal that membrane anchoring of the inhibitor—whether through its native TMD sequence, a polyleucine helix, or the TMH of the sigma-1 receptor (S1R)—is necessary for full inhibition (Extended Fig. 5j). These data, consistent with a previous study^62^, suggest that while the functional segment lies in the luminal domain, the TM helix enhances potency by localizing the inhibitor to the ER membrane.

## Discussion

The five known viral TAP inhibitors, encoded by members of the herpesvirus and poxvirus families, provide a striking example of functional convergent evolution^5^. Despite having no sequence or structural homology, these proteins have independently acquired the ability to block antigen presentation via TAP^5^. The structures presented in this study compile a structural atlas of viral TAP inhibition and reveal the molecular basis by which similar selective pressures can give rise to convergent biochemical solutions.

Consistent with earlier biochemical analyses^33,47,52,53^, the structures show that viral TAP inhibitors arrest the transporter at different stages of its transport cycle. ICP47 from HSV and BNLF2a from EBV engage the inward-facing, NBD-separated “resting” state, occluding the cytosolic vestibule and thereby preventing both peptide binding and NBD dimerization. By contrast, hUS6 from hCMV, CPXV012 from cowpox virus, and bUL49.5 from bovine herpesvirus 1 capture TAP in the outward-facing, NBD-dimerized conformation that normally supports peptide release into the endoplasmic reticulum. These inhibitors insert into the ER-luminal cavity and stabilize outward-facing state, preventing its reset for subsequent rounds of peptide translocation. All inhibitors except for ICP47 benefit from ER anchoring by a TM helix. Membrane tethering increases the local concentration of the inhibitors on the ER membrane, and in the case of bUL49.5, connects the protein with a cytoplasmic degradation module.

Although they target opposite ends of the transport cycle, all five proteins converge on a common mechanism (Fig. 6): by locking TAP in a single conformation, they halt its alternating-access cycle and abolish peptide translocation. This blockade enables the viruses to evade cytotoxic-T-lymphocyte surveillance and establish lifelong persistence in the host. Because TAP is highly conserved across vertebrates, zoonotic viruses that encode TAP inhibitors, such as rhUS6, bUL49.5, and CPXV012, either already inhibit human TAP or would require only minor adaptations to do so. These findings have important implications for viral evolution and the zoonotic potential of immune-evasive pathogens.

**Fig. 6.**
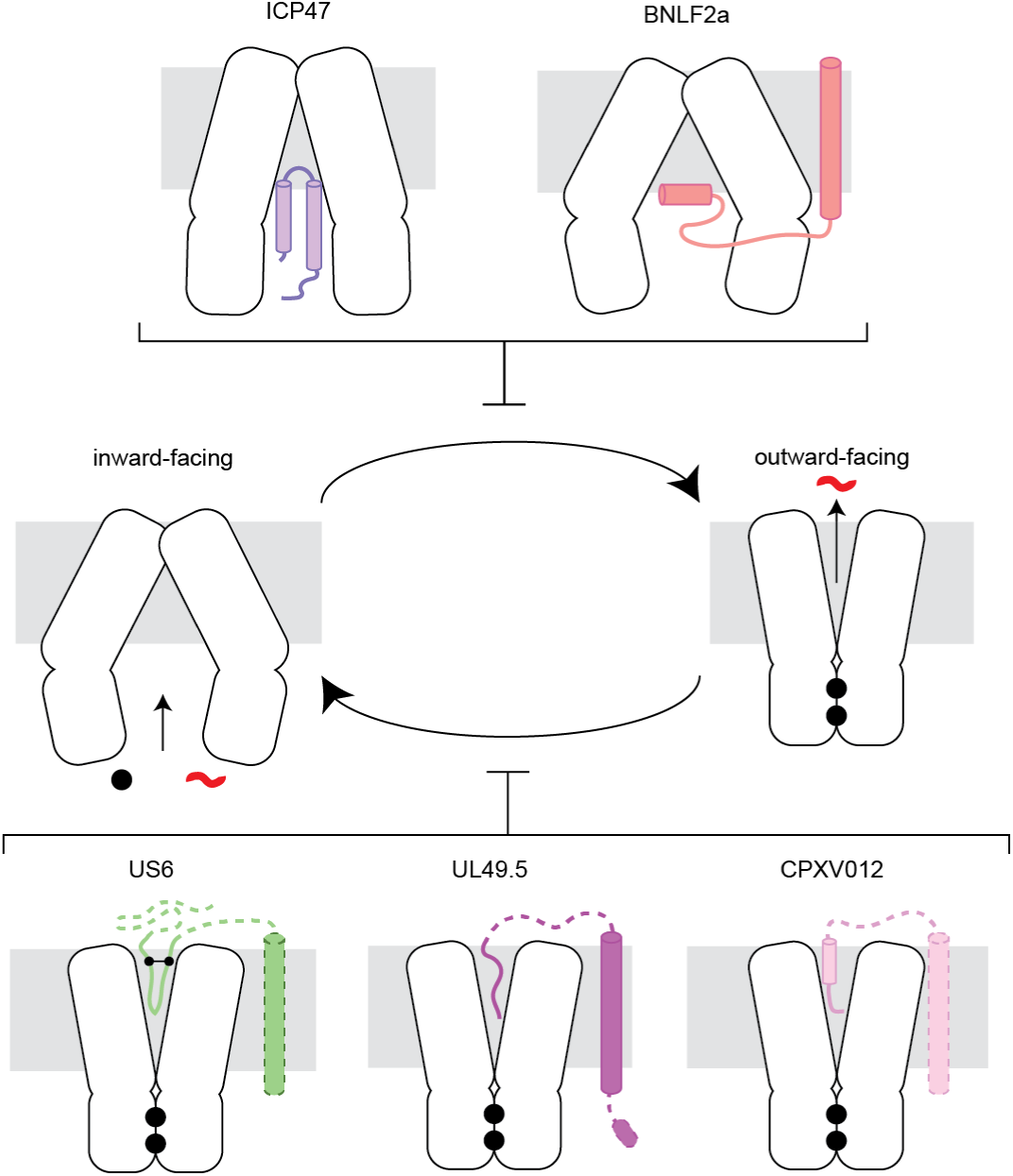
Diverse viral inhibitors stalling TAP from alternating-access transition. Helices are shown as cylinders. Dotted lines indicate regions not observed in the cryo-EM density. Nucleotide and the peptide substrate are shown as black dots and red lines, respectively. The disulfide observed in US6 is marked by connected black spheres.

A recurring mechanistic theme among these viral inhibitors is their exploitation of the TAP peptide-binding site as an anchoring point. In this study, we demonstrate that the N-and C-pockets of TAP, enriched in charged residues that normally coordinate the free termini of peptide antigens, are frequently co-opted by viral proteins to enhance binding affinity. These charged surfaces are engaged either through terminal groups or via side chains of opposite charge that mimic native substrate interactions. This strategy not only sterically occludes the translocation pathway but, in many cases, also stabilizes TAP in conformations that are incompatible with transport. The repeated targeting of these substrate-recognition elements by inhibitors that are structurally and evolutionarily unrelated highlights a shared point of vulnerability in the TAP transport cycle. Understanding these recurring strategies may guide the design of immunomodulatory therapeutics that mimic or disrupt these viral mechanisms to either suppress autoimmunity or enhance antigen presentation in cancer and infectious disease.

## Methods

### Cell Culture

*Spodoptera frugiperda* Sf9 cells (ATCC CRL-1711) were cultured in Sf-900 II SFM medium (Gibco) and 1% (v/v) antibiotic-antimycotic (Gibco) at 27°C. HEK293S GnTI-cells (ATCC CRL-3022) were cultured in Freestyle 293 medium (Gibco) supplemented with 2% (v/v) FBS and 1% (v/v) antibiotic-antimycotic at 37°C with 8% CO_2_ and 80% humidity. HEK293T cells (ATCC CRL-3216) were cultured in Dulbecco’s DMEM (Gibco) supplemented with 10% (v/v/) FBS, 1% GlutaMax (GIBCO), and 1% (v/v) antibiotic-antimycotic at 37°C with 8% CO_2_ and 80% humidity. TAP knockout (KO) HEK293S GnTI-cells were generated as previously described^49^. All commercial cell lines were authenticated by their respective suppliers. Cell lines were tested monthly for mycoplasma contamination by PCR using a Universal Mycoplasma Detection Kit (ATCC) and verified to be negative.

### Cloning

Human TAP1 with a C-terminal SpyCatcher and TAP2 were cloned into separate BacMam baculovirus expression vectors to generate pEG TAP1-SpyCatcher and pEG TAP2, respectively. For TAP1/TAP2 complex coexpression, the individual plasmids were combined via SphI and AvrII restriction sites in pEG TAP1-SpyCatcher and via SphI and NheI restriction sites in pEG TAP2 to generate pEG TAP1-SpyCatcher/TAP2. The catalytic glutamate in the consensus ATPase site of TAP2 was mutated to a glutamine to generate TAP(EQ). DNA encoding the viral inhibitors were synthesized (Genscript) and cloned into a BacMam vector containing an N-or C-terminal SpyTag and PreScission Protease-cleavable GFP tag as shown in Extended Fig. 1a.

SpyCatcher was amplified from SpyCatcher003, which was a gift from Mark Howarth (Addgene plasmid # 133447; http://n2t.net/addgene:133447; RRID:Addgene_133447). For the lentiviral constructs, DNA encoding the viral inhibitors were cloned into separate FM5 expression vectors with a C-terminal GFP linked by an internal ribosome entry site (IRES). All constructs were validated by Sanger sequencing.

Sequences for the viral proteins were codon optimized based on the protein sequences retrieved from UniProt using accession numbers P0C739 (BNLF2a), P14334 (hUS6), Q2FAB2 (rhUS6), Q77CE4 (bUL49.5), and Q8QN49 (CPXV012). BNLF2a ΔTM lacks residues D43-I60. BNLF2a pL-TM and VAMP-TM has this region substituted with polyleucine or the transmembrane helix of VAMP2 (M95-F114; UniProt accession: P63027), respectively. hUS6 ΔTMH lacks residues F145-S183 while hUS6 pL-TM has this region substituted with polyleucine. rhUS6 ΔTMH lacks residues G143-H170S while rhUS6 pL-TM has this region substituted with polyleucine. CPXV012 ΔTM has F2-I27 replaced with the signal sequence of US6. CPXV012 pL-TM and S1R-TM has S7-I27 substituted with polyleucine or the transmembrane helix of the sigma 1 receptor (A10-L30; UniProt accession: Q99720), respectively.

### Baculovirus production

Bacmid carrying TAP was generated by transforming DH10Bac *E. coli* cells with the pEG plasmids containing the gene of interest. Recombinant baculovirus was generated by transfecting Sf9 cells with bacmid using Cellfectin II (Invitrogen). Five days after transfection, P1 baculoviruses were harvested from Sf9 cell media by filtering through a 0.22 µm filter. Baculoviruses were amplified two more times to generate P2 and P3 before using for cell transduction.

### Lentivirus production

Lentivirus preparations were generated by transfecting HEK293T cells with a master mix of the FM5 plasmid containing the gene of interest, VSVG, and psPAX2 packaging plasmids at a 1:1:3 (w/w) ratio using Lipofectamine 3000 (Invitrogen). Virus was harvested from the media 36 hours after transfection, clarified by centrifugation for 5 min at 500 x g, and either used immediately or stored in aliquots at-80°C.

### Analysis of whole cell lysates

HEK293S GnTI-cells were grown in a 6-well plate and infected with 5% (v/v) P3 baculovirus at 37°C for 36 hours. Cells were harvested by resuspension in 1 ml of buffer containing 50 mM HEPES (pH 8.0 with KOH) and 150 mM KCl and spun down in 1.5 ml tubes for 5 min at 4,000g at 4°C. The cell pellets were resuspended in 1 ml of the same buffer supplemented with 1% GDN and incubated for 60 min at 4°C. Cell lysates were clarified by centrifugation at 20,000g for 2 x 30 min at 4°C. Supernatants were immediately used for analysis by fluorescent size exclusion chromatography (FSEC) or SDS-PAGE. FSEC analyses were performed using a Superose 6 10/300 column (GE Healthcare) pre-equilibrated with SEC buffer. SDS-PAGE analyses were performed at 180V for 75 min using precast 4-20% Tris-HCl polyacrylamide gradient gels (ThermoFisher). In-gel fluorescence was detected using a ChemiDoc MP imaging system (Bio-Rad).

### Flow Cytometry

MHC-I surface expression was analyzed using an allophycocyanin (APC)-coupled antibody W6/32 (eBioscience), which recognizes an epitope shared among all HLA-A,B,C molecules. HEK293S GnTI-cells grown in a 96-well plate were infected with 5% (v/v) P1 baculovirus (for experiments presented in Fig. 1 and Extended Fig. 1) or 10% (v/v) lentivirus (for experiments presented in Fig. 2, 3, 4) or transiently transfected (for experiments presented in Fig. 5) at 37°C for 36 hours. Cells were resuspended in 100 µl FACS blocking buffer (phosphate buffered saline (PBS) supplemented with 5% (w/v) bovine serum albumin (BSA) (Sigma) and centrifuged at 400g for 5 min at 25°C. Cells were resuspended in FACS blocking buffer supplemented with antibody added at 5 µg ml^−1^ and incubated for 30 min at 4°C in the dark. Subsequently, the cells were washed two times with FACS blocking buffer. The resulting cell pellets were resuspended in FACS buffer (PBS supplemented with 0.5% (w/v) BSA and 0.1% (w/v) sodium azide and counted using an Attune NxT Flow Cytometer (ThermoFisher). Gating for live cells with moderate levels of the GFP were used to compare MHC-I expression. Data were analyzed using FlowJo. Technical replicates were measured in parallel from different cells on the same day and biological replicates were measured from different cells on different days.

### Protein expression

Proteins were expressed in HEK293S GnTI-cells infected with a baculovirus mixture containing 3% (v/v) of Spycatcher-labeled TAP and 6% (v/v) of a SpyTag-labeled viral inhibitor at a density of 2.5-3.0 x 10^6^ cells/ml. WT TAP was used for samples containing BNLF2a, hUS6, or rhUS6 and TAP(EQ) was used for samples containing bUL49.5 and CPXV012. Cells were induced with 10 mM sodium butyrate 8-12 hours after infection and cultured at 30°C for another 48 hours. Cells were harvested, snap frozen in liquid nitrogen, and stored at-80°C.

### Protein purification

Cells were thawed and resuspended in lysis buffer containing 50 mM HEPES (pH 6.5 with KOH), 400 mM KCl, 2 mM MgCl_2_, 1mM dithiothreitol (DTT), 20% (v/v) glycerol, 1 μg ml^−1^ pepstatin A, 1 μg ml^−1^ leupeptin, 1 μg ml^−1^ aprotinin, 100 μg ml^−1^ soy trypsin inhibitor, 1 mM benzamidine, 1 mM phenylmethylsulfonyl fluoride (PMSF) and 3 µg ml^−1^ DNase I. For samples containing nucleotide, all buffers were supplemented with 1 mM ATP starting from cell lysis. Cells were lysed by three passes through a high-pressure homogenizer at 15,000 psi (Emulsiflex-C3; Avestin). Unbroken cells and cell debris were removed by one low speed spin at 4000g for 15 min at 4°C. The supernatant was subjected to a second round of ultracentrifugation at 100,000 x g for 1 hour at 4°C in a Type 45Ti rotor (Beckman) to pellet cell membranes. Membranes were resuspended by manual homogenization in a dounce in lysis buffer supplemented with 1% glycol-diosgenin (GDN) (Anatrace) and incubated for 1 hour at 4°C. The insoluble fraction was removed by centrifugation at 75,000g for 30 min at 4°C and the supernatant was applied to NHS-activated Sepharose 4 Fast Flow resin (GE Healthcare) conjugated with GFP nanobody pre-equilibrated in lysis buffer. After 1 hour, the resin was washed with 10 column volumes of wash buffer containing 50 mM HEPES (pH 6.5 with KOH), 400 mM KCl, 10% glycerol, 1 mM DTT, and 0.01% GDN. To cleave off the GFP tag, PreScission Protease was added to a final concentration of 0.35 mg ml^−1^ and incubated for 12 hours at 4°C. The cleaved protein was eluted with 5 column volumes of wash buffer and collected by passing through a Glutathione Sepharose 4B resin (Cytiva) to remove the PreScission Protease. The eluate was then concentrated using a 15 ml Amicon spin concentrator with a 100-kDa molecular weight cutoff membrane (Millipore) and purified by size exclusion chromatography (SEC) using a Superose 6 Increase 10/300 column (GE Healthcare) pre-equilibrated with SEC buffer containing 50 mM HEPES (pH 6.5 with KOH), 200 mM KCl, 1 mM DTT and 0.004% GDN. Peak fractions were pooled using a 4 ml Amicon spin concentrator with a 100-kDa molecular weight cutoff membrane (Millipore) and used immediately for grid preparation.

### Cryo-EM sample preparation and imaging

TAP purified from gel filtration was concentrated to ∼6 mg ml^−1^. An additional 5 mM of ATP was applied to each sample immediately before freezing except for the sample containing BNLF2a. Grids were prepared by applying 3.5 µL of protein onto a glow discharged Quantifoil R0.6/1.0 400 mesh holey carbon Au grid with no wait time. The grids were blotted for 3 sec and plunged frozen into liquid ethane using an FEI Mark IV Vitrobot at 6°C and 100% humidity.

All the cryo-EM data were collected using a 300 kV Titan Krios transmission electron microscope equipped with a Gatan K3 camera except for the bUL49.5 sample, which was collected with a Thermo Scientific Falcon 4i camera. All micrographs were collected using SerialEM^63^ in super-resolution mode except for the bUL49.5 sample. Data collection parameters for different samples are summarized in Extended Data Table 1.

### Cryo-EM Image Processing

For the BNLF2a, US6, and RH185 datasets, super-resolution image stacks were gain-normalized, binned by 2, and corrected for beam-induced motion using MotionCor2^64^. Contrast transfer function (CTF) parameters were estimated using CTFFIND4^65^.

For the BNLF2a dataset, particles were auto-picked from the motion-corrected micrographs with crYOLO using its general model^66^, extracted in RELION^67^, and imported into cryoSPARC^68^. The picked particles were subjected to multiple rounds of 2D classification, and the resulting particles were subjected to *ab initio* reconstruction with five classes. Three classes resembled an empty micelle while the other two classes resembled TAP, but with varying continuous density for the NBDs. To improve the density of the NBDs, all the particles from 2D classification were subjected to iterative rounds of heterogenous refinement using the two reconstructions that resembled TAP from the *ab initio* as input models. The resulting particles that gave reconstructions with complete NBDs were then subjected to tandem non-uniform refinement followed by local refinement with a protein mask excluding the micelle. These particles were imported into RELION using the csparc2star.py script^69^ and subjected to Bayesian particle polishing^70^ and non-uniformed refined to obtain the final map.

For the hUS6 dataset, particles were auto-picked from the motion-corrected micrographs with crYOLO using its general model, extracted in RELION, imported into cryoSPARC, and subjected to 2D classification. The best classes with clear TAP features were selected to train a Topaz^71^ model that was used for another round of particle picking. After multiple rounds of 2D classification, the resulting particles were pooled and duplicate picks were removed. The resulting particle stack was subjected to *ab initio* reconstruction with 3 classes with one class resembling NBD-dimerized TAP. The particle stack was further sorted by multiple rounds of heterogeneous refinement and one round of non-uniform refinement. This reconstruction consisted of 75,164 particles with clear TAP density and visible, but broken hUS6 density. These particles were imported into RELION, subjected to Bayesian particle polishing, and imported back into cryoSPARC for another round of heterogeneous and non-uniform refinement. The particles were imported back into RELION, CTF refined and 3D classified without alignment using a mask excluding the micelle. One class had well defined transmembrane helices and continuous US6 density that was then used for non-uniform refinement to obtain the final map.

For the rhUS6 dataset, particles were auto-picked from a subset of a thousand micrographs using the Laplacian-of-Gaussian filter, extracted in RELION, imported into cryoSPARC, and subjected to 2D classification. The best classes with clear TAP features were selected to train a Topaz model that was used to pick particles from all the micrographs. After multiple rounds of 2D classification, an initial model was generated by *ab initio* reconstruction with 3 classes. One class showed clear density for NBD-dimerized TAP and this model was used for further sorting by multiple rounds of heterogeneous refinement with 2 junk classes. Non-uniform refinement of the resulting 252,941 particles had clear density for the rhUS6 transmembrane helix and loop, but broken density for the linker between the two. 3D classification in RELION without alignment with a mask excluding the micelle was performed to identify a class with the most distinct TAP and rhUS6 density. To improve the quality of the rhUS6 density, CTF refinement and another round of 3D classification without alignment with a mask around the rhUS6 density was performed. One class exhibited clear rhUS6 density. This final particle stack was then subject to CTF refinement, non-uniform refinement, Bayesian particle polishing, and one more round of non-uniform refinement to generate the final map.

For the bUL49.5 dataset, all initial data processing was carried out in cryoSPARC. Raw movies in EER format were divided into 52 fractions and were corrected for beam-induced motion using the Patch motion correction tool. CTF parameters were estimated using the Patch CTF estimation tool. 2,741,436 were auto-picked using the Blob picker tool, extracted, and sorted by multiple rounds of 2D classification. *Ab initio* reconstruction with three classes yielded one class with clear NBD-dimerized TAP features and this model was used for further sorting by multiple rounds of heterogeneous refinement with 2 junk classes. Non-uniform refinement of the resulting 965,440 particles yielded a reconstruction with good TAP density, but weak density for the bUL49.5 transmembrane helix and amorphous density within the TAP translocation pathway. Further 3D classification without alignment with a mask excluding the micelle in RELION was used to sort out particles with the strongest TAP density. Non-uniform refinement of these 303,714 particles yielded two major conformations defined by a major movement in TAP2 TM3 and in the position of inhibitor density. Bayesian particle polishing of this particle stack and then subsequent 3D classification without alignment with a mask on inhibitor density and TAP transmembrane helices was then used to sort out the kinked and unkinked conformations of TAP.

For the CPXV012 dataset, all initial data processing was carried out in cryoSPARC. Super-resolution image stacks were gain-normalized, binned by 2, and corrected for beam-induced motion using the Patch motion correction tool. CTF parameters were estimated using the Patch CTF estimation tool. Particle were auto-picked using the Blob picker tool, extracted, and sorted by multiple rounds of 2D classification. *Ab initio* reconstruction with three classes yielded one class with clear density for transmembrane helices that was then non-uniform refined. This stack of 309,154 particles was subject to Bayesian particle polishing and another round of 3D classification without alignment with a mask excluding the micelle. The class with the clearest density in the transmembrane helices was selected for another round of 3D classification without alignment. The best class with the clearest density for CPXV012 was then selected for one more round of non-uniform refinement to generate the final map.

FSC curves were generated in cryoSPARC, and resolutions were reported based on the 0.143 criterion. Masking and B-factor sharpening were determined automatically in cryoSPARC during refinement.

### Model Building and Refinement

The sharpened and unsharpened maps from local refinement were used for model building. Molecular models of TAP were initially built based on the cryo-EM structures of apo TAP in the inward-facing state and TAP(EQ) in the outward-facing state. Models for the viral inhibitors were generated using ModelAngelo. The models were then docked into the density, iteratively edited and refined in Coot^72^, ISOLDE^73^, and PHENIX^74^. The quality of the final models were evaluated by MolProbity^75^. Refinement statistics are summarized in Extended Data Table 1.

### Figure preparation

Cryo-EM maps and atomic models depictions were generated using UCSF ChimeraX^76^. Maps colored by local resolution were generated using cryoSPARC. Multiple sequence alignments were generated using Clustal Omega^77^. Graphs and associated statistics were prepared using GraphPad Prism 9. Structural biology software used in this project was managed by SBGrid^78^. All Figures were prepared using Adobe Illustrator.

## Data availability

Cryo-EM density maps have deposited in the Electron Microscopy Data Bank under accession codes EMD-70314, EMD-70315, EMD-70316, EMD-70137, EMD-70241, and EMD-70246. The corresponding atomic models have been deposited in the Protein Data Bank under the accession codes 9OCG, 90CH, 90CI, 9OCJ, 9O94, and 909D. The raw micrographs for all datasets have been deposited into the Electron Microscopy Public Image archive under the accession codes EMPIAR-12714, EMPIAR-12759, EMPIAR-12760, EMPIAR-12761, and EMPIAR-12762. Any additional data reported in this paper is available from the lead contact upon request.

## Acknowledgements

We thank Rui Yan and Zhiheng Yu at the Howard Hughes Medical Institute (HHMI) Janelia Cryo-EM Facility and Mark Ebrahim, Johanna Sotiris, and Honkit Ng at the Evelyn Gruss Lipper Cryo-EM Resource Center at Rockefeller University for assistance in electron microscopy data collection; and members of the Chen and MacKinnon laboratories for helpful discussions. J.L. is a HHMI Fellow of the Helen Hay Whitney Foundation, and J.C. is an investigator of the HHMI. V. M. is supported by a Medical Scientist Training Program grant from the National Institute of General Medical Sciences of the National Institutes of Health (T32GM007739) to the Weill Cornell/Rockefeller/Sloan Kettering Tri-Institutional MD-PhD Program. This research was also supported by the Stavros Niarchos Foundation (SNF) as part of its grant to the SNF Institute for Global Infectious Disease Research at The Rockefeller University.

## Contributions

J.L., V.M., and J.C. designed experiments; J.L. and V.M. performed experiments; J.L., V.M., and J.C. analyzed data and wrote the paper.

**Corresponding authors:** Correspondence to Jue Chen.

