## Supplemental Information for "A Structural Atlas of TAP Inhibition by Herpesviruses and Poxviruses"

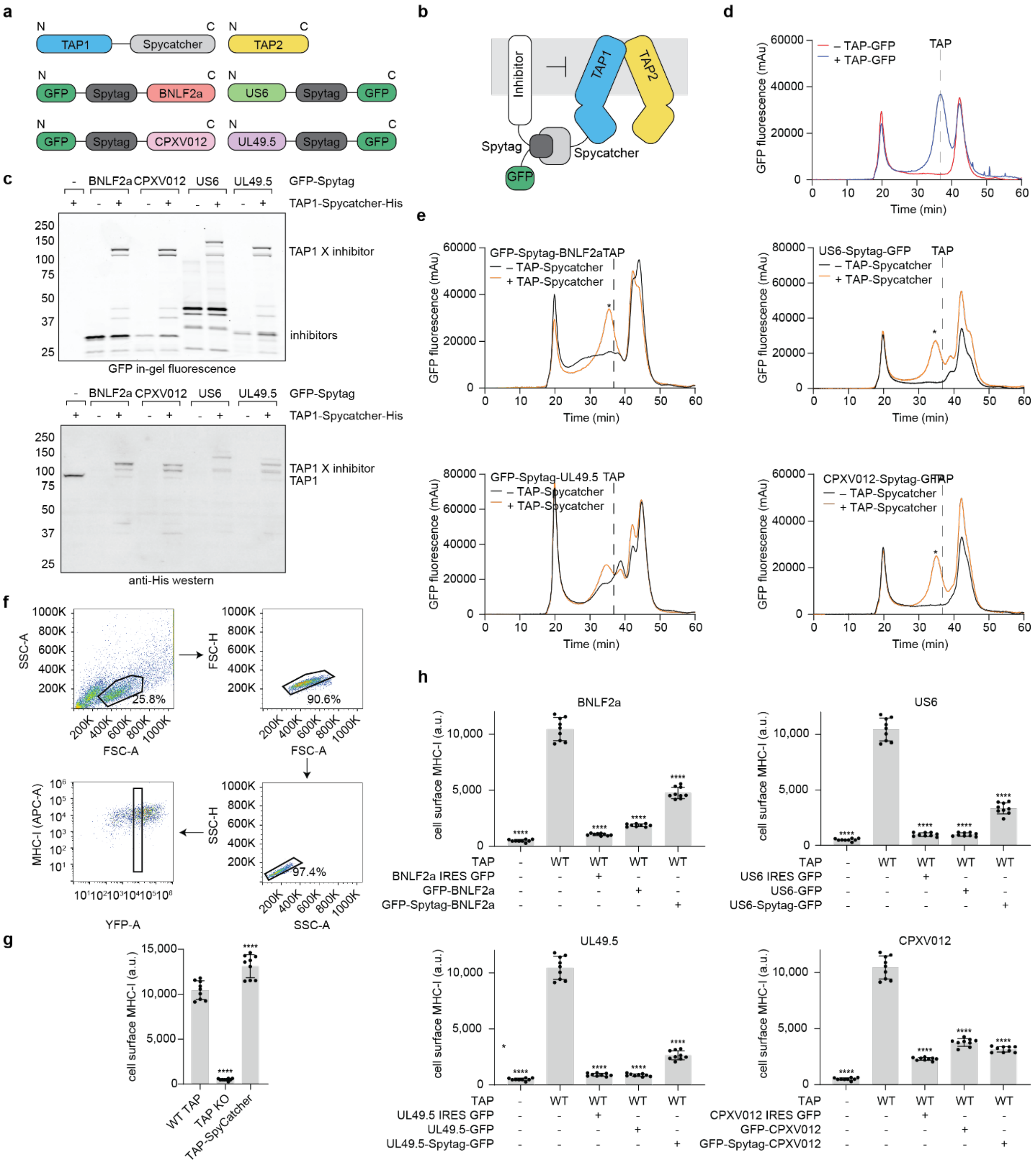


**Extended Fig. 1. Capture of virally inhibited TAP complexes using the SpyCatcher-SpyTag system.** **a.** Schematic of the individual SpyCatcher- and SpyTag-labeled constructs. **b.** Schematic of the general SpyCatcher-SpyTag labeling strategy. **c.** The TAP/inhibitor complexes form efficiently in cells. SDS-PAGE of cell lysates expressing combinations of SpyCatcher-labeled TAP and different GFP-SpyTag-labeled viral inhibitors as visualized by in-gel fluorescence and Western Blot. **d.** TAP migrates as a monodisperse peak as judged by fluorescence size exclusion (FSEC). The peak corresponding to TAP is marked by a dashed line. **e.** The TAP/inhibitor complexes migrate as a monodisperse peak. The peak corresponding to TAP and the TAP/inhibitor complex are marked by a dashed line and asterisk, respectively. **f.** Representative FACS plots and gating strategy. Data shown are from HEK293S cells expressing YFP only. **g.** The TAP-SpyCatcher construct is functional. Flow cytometry analysis of MHC-I cell surface levels in TAP knockout cells transduced with baculovirus expressing WT or SpyCatcher-tagged TAP. Data represent the mean fluorescence intensity and standard error of three technical replicates of three biological replicates (n=9). Error bars represent the SEM. Statistical significance relative to the TAP-Spycatcher sample was tested by one-way analysis of variance. ns, not significant; ****P<0.0001. **h.** The SpyTag-labeled viral inhibitors are functional. Flow cytometry analysis of MHC-I cell surface levels in TAP knockout cells transduced with baculovirus expressing combinations of TAP and the different viral inhibitors. Data represent the mean fluorescence intensity and SEM of three technical replicates of three biological replicates (n=9). Samples from TAP knockout (KO) cells and wild-type (WT) cells alone are replicated in each chart. Samples with SpyTag-GFP labelled viral inhibitors are replicated from Fig. 1b. Statistical significance relative to the wild-type was tested by one-way analysis of variance. ns, not significant; ****P<0.0001.

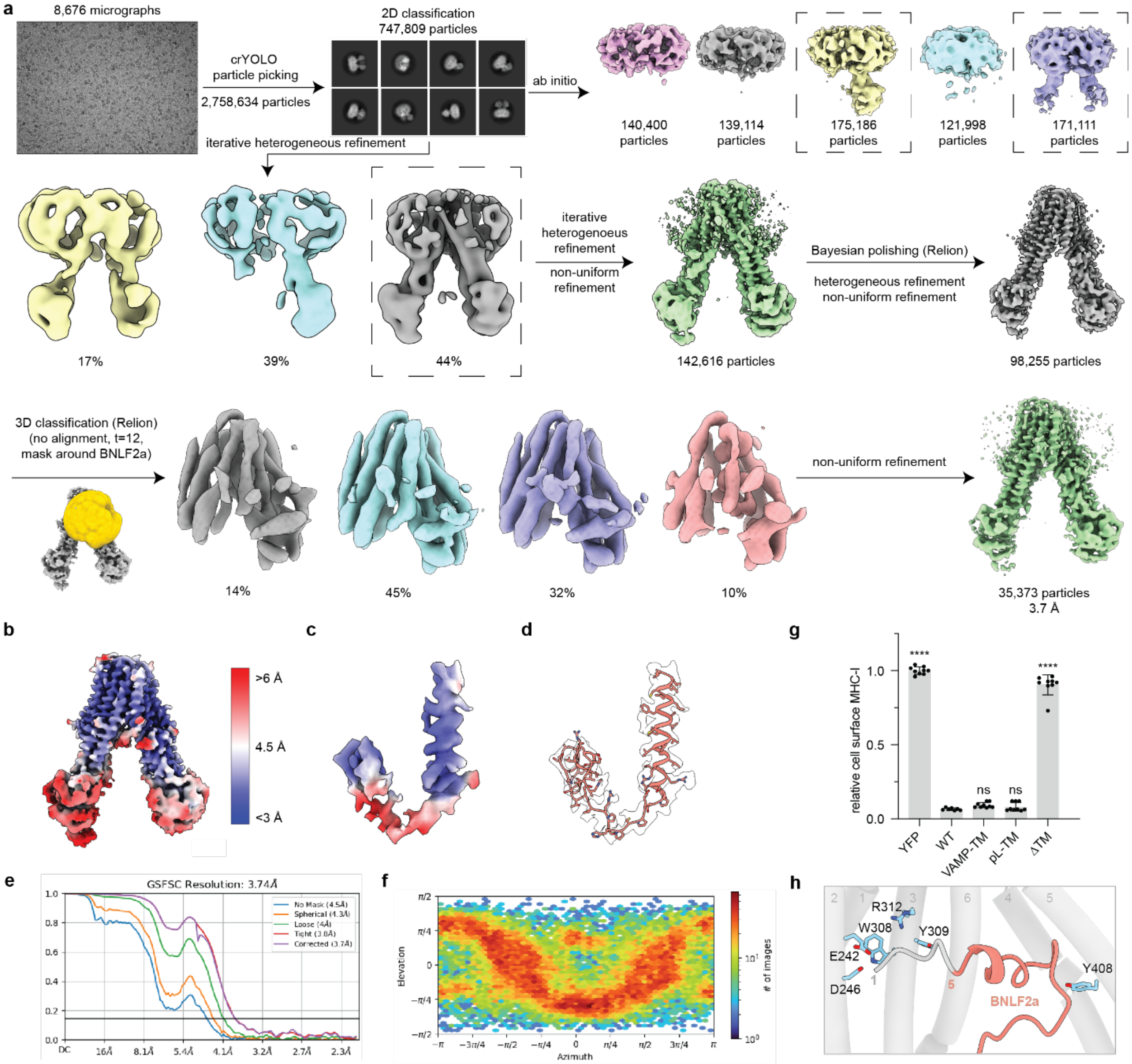


**Extended Fig. 2. Structural and biochemical analysis of the TAP/BNLF2a complex.**
**a.** Flowchart of the image processing of BNLF2a-inhibited TAP. **b.** Local resolution map of BNLF2a-inhibited TAP. **c.** Local resolution map of the BNLF2a. Reconstruction is contoured to 0.2 SDs. **d.** Map to model fit of BNLF2a. Reconstruction is contoured to 0.15 SDs. **e.** Fourier shell correlation (FSC) curves of the final reconstruction of BNLF2a-inhibited TAP. **f.** Angular distribution of the final reconstruction of BNLF2a-inhibited TAP. **g.** Flow cytometry analysis of the HEK293S cells expressing BNLF2a transmembrane helix swap variants coupled to YFP with an internal ribosomal entry site (IRES) element. Data represent the relative MHC-I cell surface levels normalized to the YFP only sample and represent three technical replicates of three biological replicates (n=9). Error bars represent the SEM. Statistical significance relative to the wild-type was tested by one-way analysis of variance. ns, not significant; ****P<0.0001. **h.** Putative binding mode of the BNLF2a N-terminus to the TAP N-pocket. Residues 1-5 of BNLF2a are not visible in the cryo-EM density and are putatively modeled.

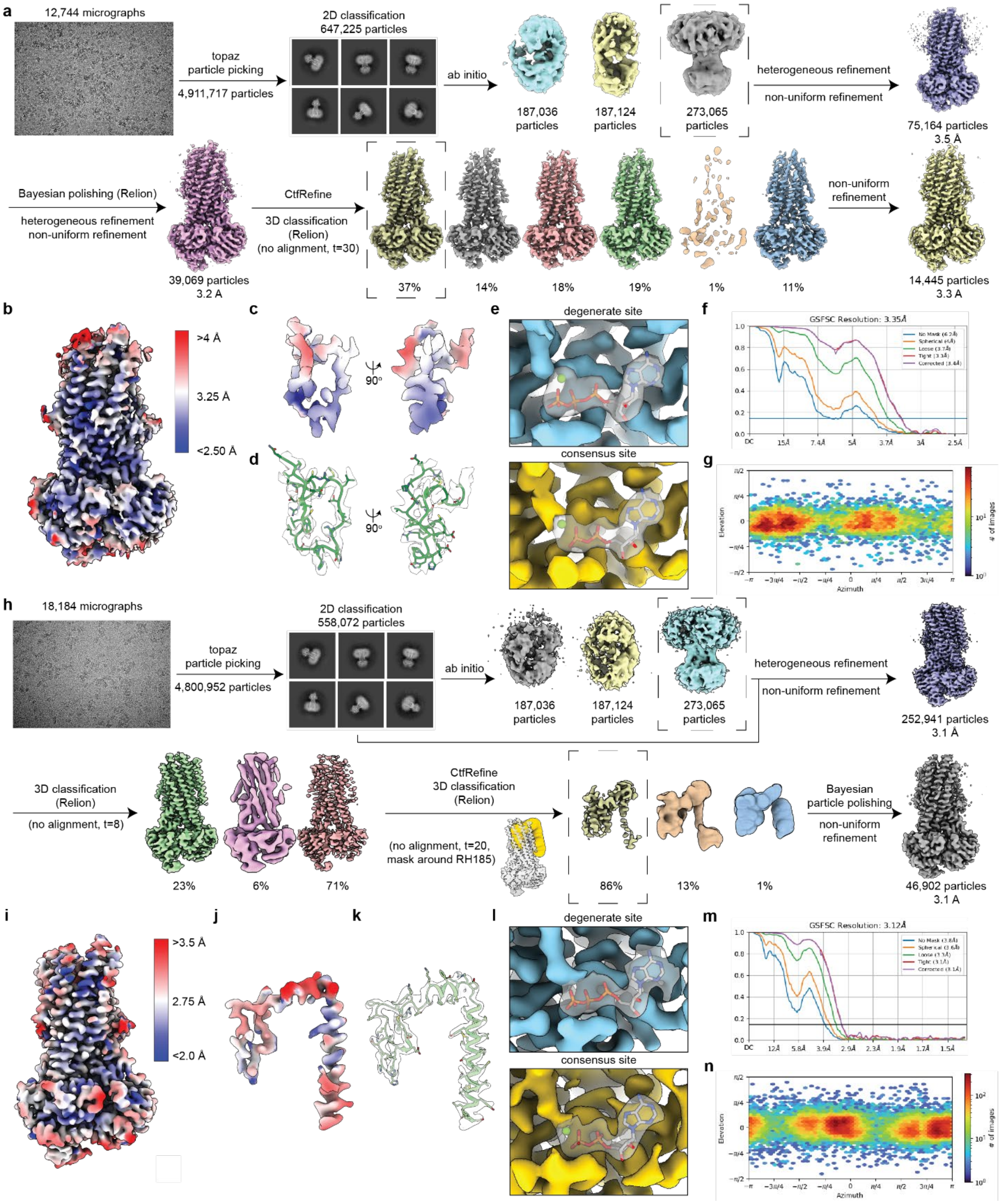


**Extended Fig. 3. Cryo-EM processing of hUS6-inhibited and rhUS6-inhibited TAP structures.**
**a.** Flowchart of the image processing of hUS6-inhibited TAP. **b.** Local resolution map of the final overall complex of hUS6-inhibited TAP. **c.** Local resolution map of hUS6. Reconstructions are contoured to 0.3 SDs. **d.** Map to model fit of hUS6. Reconstructions are contoured to 0.3 SDs. **e.** Map to model fit around

the nucleotide binding sites of TAP. Reconstructions are contoured to 0.4 SDs. **f.** Fourier shell correlation (FSC) curves of the final reconstruction of hUS6-inhibited TAP. **g.** Angular distribution of the final reconstruction of hUS6-inhibited TAP. **h.** Flowchart of the image processing of rhUS6-inhibited TAP **i.** Local resolution map of the final overall complex of rhUS6-inhibited TAP. **j.** Local resolution map. Reconstructions are contoured to 0.3 SDs. **k.** Map to model fit. **l.** Map to model fit around the nucleotide binding sites of TAP. Reconstructions are contoured to 0.3 SDs. **m.** Fourier shell correlation (FSC) curves of the final reconstruction of rhUS6-inhibited TAP. **n.** Angular distribution of the final reconstruction of rhUS6-inhibited TAP.

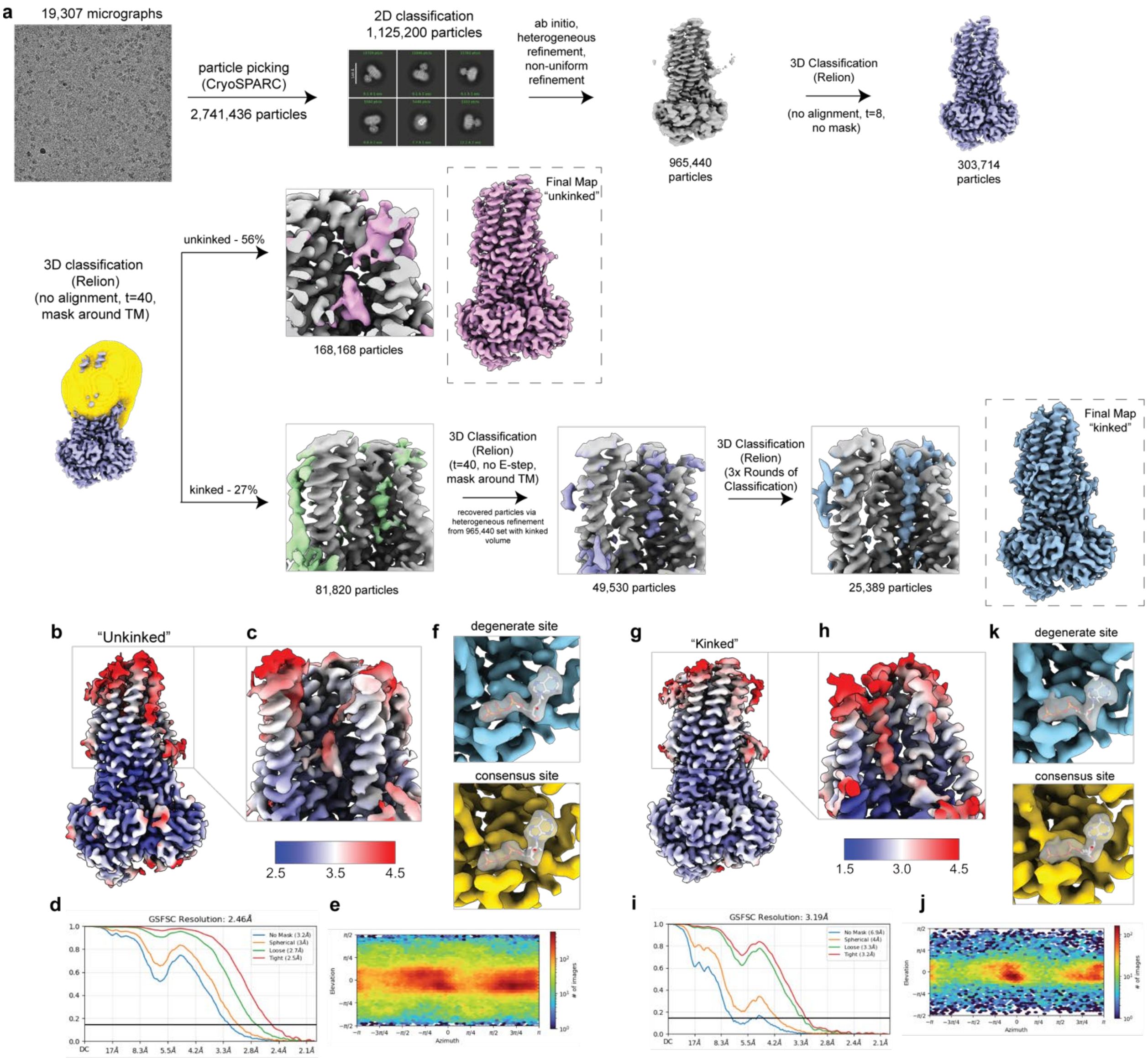


**Extended Fig. 4. Cryo-EM processing of bUL49.5-inhibited TAP(EQ).**
**a.** Flowchart of the image processing of bUL49.5-inhibited TAP. **b,c.** Local resolution map of bUL49.5-TAP(EQ) in the "unkinked" conformation. Reconstructions are contoured to 0.188 SDs. **d.** Fourier shell correlation (FSC) curves of the "unkinked". **e.** Angular distribution of the final reconstruction. **f.** Map to model fit around the nucleotide binding sites of TAP. Reconstructions are contoured to 0.188 SDs. **g,h.** Local resolution map of bUL49.5-TAP(EQ) in the "kinked" conformation. Reconstructions are contoured to 0.135 SDs. **i.** Fourier shell correlation (FSC) curves of the "kinked". **j.** Angular distribution of the final reconstruction. **k.** Map to model fit around the nucleotide binding sites of TAP. Reconstructions are contoured to 0.135 SDs.

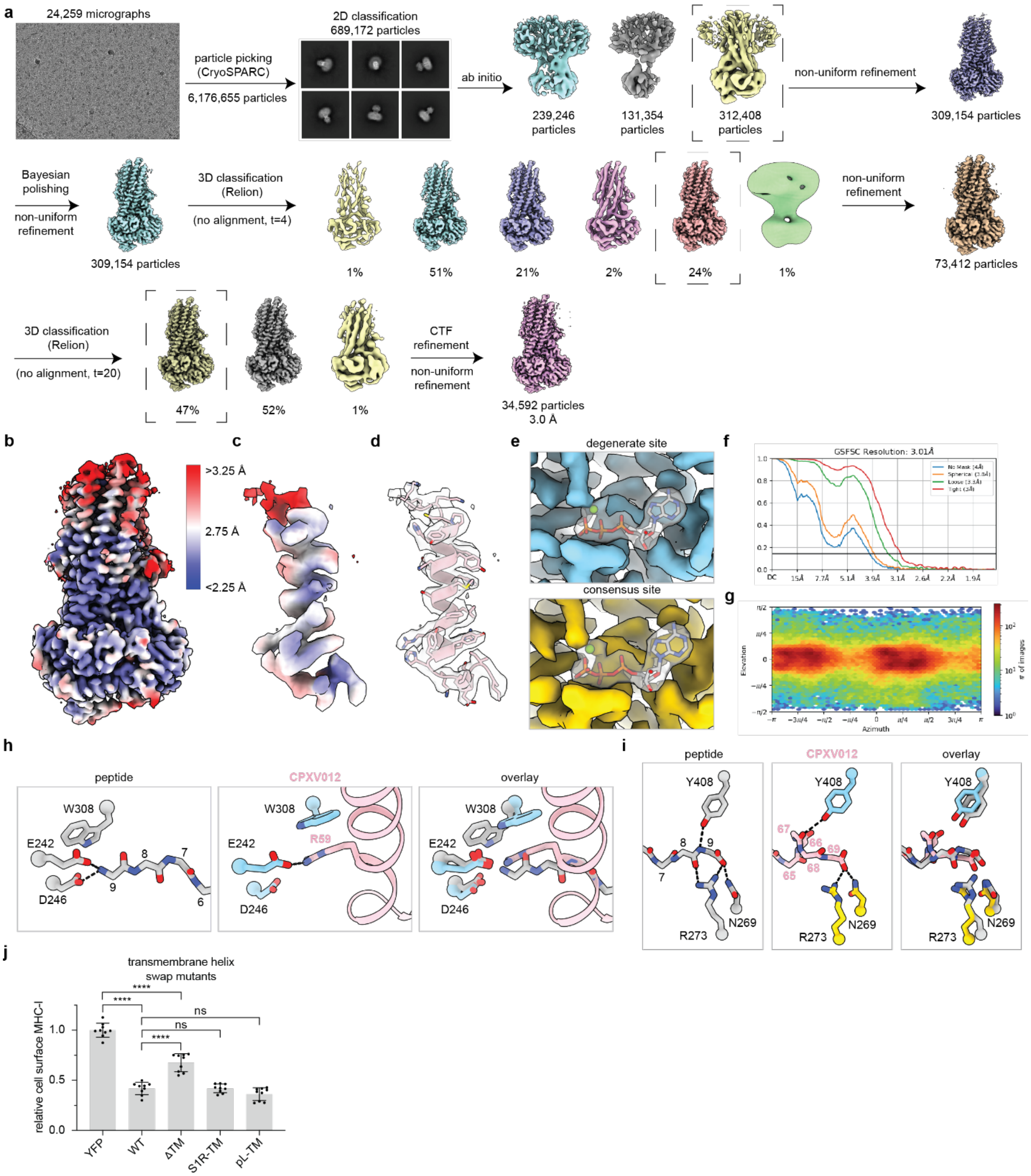


**Extended Fig. 5. Structural and biochemical analysis of CPXV012-inhibited TAP(EQ).**
**a.** Flowchart of the image processing of CPXV012-inhibited TAP(EQ). **b.** Local resolution map of the final overall complex of CPXV012-inhibited TAP(EQ). **c.** Local resolution map of CPXV012. Reconstructions are contoured to 0.16 SDs. **d.** Map to model fit of CPXV012. Reconstructions are contoured to 0.16 SDs. **e.** Map to model fit around the nucleotide binding sites of TAP. Reconstructions are contoured to 0.3 SDs. **f.** Fourier shell correlation (FSC) curves of the final reconstruction of CPXV012-inhibited TAP(EQ). **g.** Angular distribution of the final reconstruction of CPXV012-inhibited

TAP(EQ). **h.** Comparison of peptide (PDB:8T4F) or CPXV012 binding to the TAP N-pocket in the inward-facing state and outward-facing state, respectively. The sidechains of the peptide are hidden for clarity. **i.** Comparison of peptide (PDB:8T4F) or CPXV012 binding to the TAP C-pocket in the inward-facing state and outward-facing state, respectively. The sidechains of the peptide are hidden for clarity. **j.** The transmembrane helix of CPXV012 is important for localizing the soluble domains to TAP. Flow cytometry analysis of HEK293S cells expressing CPXV012 transmembrane helix variants coupled to YFP with an IRES element. Data represent the relative MHC-I cell surface levels normalized to the YFP only sample and represent three technical replicates of three biological replicates (n=9). Error bars represent the SEM. Statistical significance relative to treatment with wild-type CPXV012 IRES YFP was tested by one-way analysis of variance. ns, not significant; ****P<0.0001.

**Extended Data Table 1.** Cryo-EM data collection, refinement, and validation statistics

| | TAP BNLF2a EMD-70314 PDB 9OCG | TAP hUS6 EMD-70315 PDB 9OCH | TAP rhUS6 EMD-70316 PDB 9OCI | TAP(EQ) bUL49.5 unkinked EMD-70241 PDB 9O94 | TAP(EQ) bUL49.5 kinked EMD-70246 PDB 9O9D | TAP(EQ) CPXV012 EMD-70317 PDB 90CJ |
|---|---|---|---|---|---|---|
| **Data collection and processing** | | | | | | |
| Magnification | 81,000 | 130,000 | 130,000 | 130,000 | | 105,000 |
| Voltage (kV) | 300 | 300 | 300 | 300 | | 300 |
| Electron exposure ($e^{-}$/Å$^2$) | 51 | 50 | 50 | 50 | | 45 |
| Defocus range (μm) | 0.8 to 2.5 | 0.8 to 2.0 | 0.8 to 1.8 | 0.8 to 1.8 | | 0.8 to 1.8 |
| Pixel size (Å) | 1.08 | 0.663 | 0.663 | 0.743 | | 0.86 |
| Symmetry imposed | C1 | C1 | C1 | C1 | | C1 |
| Initial particle images (no.) | 2,758,634 | 4,911,717 | 4,800,952 | 2,741,436 | | 6,176,655 |
| Final particle images (no.) | 35,373 | 14,445 | 46,902 | 168,168 | 25,389 | 34,592 |
| Map resolution (Å) | 3.7 | 3.3 | 3.1 | 2.5 | 3.2 | 3.0 |
| FSC threshold | 0.143 | 0.143 | 0.143 | 0.143 | 0.143 | 0.143 |
| **Refinement** | | | | | | |
| Initial model used (PDB) | 8T46 | 9N63 | 9N63 | 9N63 | 9N63 | 9N63 |
| Model resolution (Å) | 4.1 | 3.5 | 3.3 | 2.6 | 3.1 | 3.2 |
| FSC threshold | 0.5 | 0.5 | 0.5 | 0.5 | 0.5 | 0.5 |
| Map sharpening *B* factor (Å$^2$) | 101.8 | 48 | 67.7 | 54.6 | 30.8 | 55.8 |
| Model composition | | | | | | |
| Non-hydrogen atoms | 7777 | 8896 | 9178 | 8686 | 8367 | 9006 |
| Protein residues | 1164 | 1134 | 1170 | 1140 | 1105 | 1143 |
| Ligands | 0 | 4 | 4 | 4 | 4 | 4 |
| *B* factors (Å$^2$) | | | | | | |
| Protein (mean) | 145.62 | 120.42 | 94.73 | 83.01 | 82.81 | 94.91 |
| Ligands (mean) | N/A | 114.02 | 85.17 | 61.93 | 71.09 | 68.46 |
| R.m.s. deviations | | | | | | |
| Bond lengths (Å) | 0.011 | 0.003 | 0.003 | 0.003 | 0.004 | 0.003 |
| Bond angles (°) | 1.648 | 0.696 | 0.679 | 0.614 | 0.741 | 0.626 |
| Validation | | | | | | |
| MolProbity score | 1.20 | 1.17 | 1.17 | 1.03 | 0.89 | 1.15 |
| Clashscore | 2.20 | 3.86 | 3.84 | 2.43 | 1.51 | 3.59 |
| Poor rotamers (%) | 0.16 | 0.00 | 0.10 | 0.55 | 0.47 | 0.00 |
| Ramachandran plot | | | | | | |
| Favored (%) | 96.71 | 98.04 | 98.45 | 99.12 | 99.36 | 98.33 |
| Allowed (%) | 3.29 | 1.96 | 1.55 | 0.88 | 0.64 | 1.67 |
| Disallowed (%) | 0.00 | 0.00 | 0.00 | 0.00 | 0.00 | 0.00 |
